# Opa1 and Drp1 reciprocally regulate cristae morphology, ETC function, and NAD^+^ regeneration in KRas-mutant lung adenocarcinoma

**DOI:** 10.1101/2022.06.01.494337

**Authors:** Dane T. Sessions, Kee-Beom Kim, Jennifer A. Kashatus, Nikolas Churchill, Kwon-Sik Park, Marty W. Mayo, Hiromi Sesaki, David F. Kashatus

## Abstract

Oncogenic KRas activates mitochondrial fission through Erk-mediated phosphorylation of the mitochondrial fission GTPase Drp1. Drp1 deletion inhibits tumorigenesis of KRas-driven pancreatic cancer, but the role of mitochondrial dynamics in other Ras-driven malignancies is poorly defined. Here we demonstrate that *in vitro* and *in vivo* growth of KRas-driven lung adenocarcinoma is unaffected by deletion of Drp1, but inhibited by deletion of Opa1, the GTPase that regulates inner membrane fusion and promotes proper cristae morphology. Mechanistically, Opa1 knockout induces loss of electron transport chain (ETC) complex I assembly and activity that inhibits tumor cell proliferation through loss of NAD^+^ regeneration. Simultaneous inactivation of Drp1 and Opa1 restores cristae morphology, complex I activity and cell proliferation indicating that mitochondrial fission activity drives ETC dysfunction induced by Opa1 knockout. Our results support a model in which mitochondrial fission events disrupt cristae structure and tumor cells with hyperactive fission activity require Opa1 activity to maintain ETC function.

## Introduction

Approximately one-third of all human tumors harbor mutations in the *RAS* family of GTPases. Activating *RAS* mutations rewire cellular metabolism (Racker, Resnick and Feldman, 1985; Chiaradonna *et al*., 2006; Gaglio *et al*., 2011) and promote cell survival and proliferation. Mitochondria are critical organelles that regulate a number of processes disrupted in cancer, including ATP synthesis, redox homeostasis and programmed cell death. Mitochondria are highly dynamic and undergo opposing cycles of fusion and fission, which are regulated by four large dynamin-related GTPases. Drp1 executes mitochondrial fission whereas Mfn1/2 and Opa1 regulate fusion of the mitochondrial outer and inner membranes, respectively. The fission and fusion GTPases also contribute to ultrastructural remodeling of the mitochondrial outer and inner membranes. For example, Drp1 remodels cristae during apoptosis (Otera *et al*., 2016) and Opa1 is critical to maintain cristae fidelity (Cogliati *et al*., 2013) to support oxidative phosphorylation (OXPHOS) (Del Dotto *et al*., 2017; Quintana-Cabrera *et al*., 2018; Cretin *et al*., 2021) and resistance to apoptosis (Olichon *et al*., 2003; Frezza *et al*., 2006) through sequestration of cytochrome C (Yamaguchi *et al*., 2008).

Oncogenic signaling pathways impact mitochondrial shape in a number of ways through changing the activity of the mitochondrial dynamics machinery. For example, oncogenic KRas activates mitochondrial fission through Erk2-mediated phosphorylation of Drp1 (Kashatus *et al*., 2015; Serasinghe *et al*., 2015) and inhibition of mitochondrial fission inhibits KRas-driven glycolytic flux, cellular transformation, and pancreatic tumor growth (Nagdas *et al*., 2019). To determine the extent to which Erk-driven mitochondrial fission contributes to tumor growth in other Ras-driven malignancies, we explored the effects of mitochondrial dynamics disruption in KRas-driven lung adenocarcinoma (LUAD). Lung adenocarcinoma has the highest mortality of human cancers and 30% of LUAD tumors harbor oncogenic *KRAS* mutations; however, the role of mitochondrial dynamics in this malignancy is poorly defined compared to other KRas-driven tumors. Tissue of origin can influence the cellular metabolism of tumors with identical driver mutations (Yuneva *et al*., 2012; Mayers *et al*., 2016). Since PDAC and LUAD exhibit distinct metabolic phenotypes (Reske *et al*., 1997; Ying *et al*., 2012; Bryant *et al*., 2014; Scafoglio *et al*., 2015; Hensley *et al*., 2016; Momcilovic *et al*., 2019), we explored whether LUAD demonstrates unique sensitivities to inhibition of the mitochondrial dynamics machinery. Surprisingly, deletion of Opa1, but not Drp1, inhibits *in vitro* colony formation and ETC function in KRas-driven LUAD cells, and blocks tumor development in a *KRAS^G12D/+^; TP53^-/-^* (KP) genetically-engineered mouse model (GEMM) of LUAD. Further, Drp1 deletion completely rescues the effects of Opa1 deletion on *in vitro* colony formation and ETC function, but not *in vivo* tumor growth. Mechanistically, Opa1 deletion impacts colony formation and tumor growth through loss of proper cristae morphology and subsequent inhibition of ETC function. Consistent with recent work (Birsoy *et al*., 2015; Sullivan *et al*., 2015; Luengo *et al*., 2021), the critical ETC function disrupted by Opa1 deletion is not mitochondrial ATP synthesis but NAD^+^ regeneration by complex I. Collectively, these data suggest that Opa1-dependent cristae remodeling plays a critical role in promoting complex I-mediated NAD^+^ regeneration in proliferative cells and other cells with high levels of mitochondrial fission activity.

## Results

### Opa1 inhibition prevents KRas*-*mutant LUAD colony formation in a Drp1-dependent manner (Figure 1)

Mitochondrial shape is maintained through a balance of fission and fusion. Persistent Ras-MAPK signaling hyperactivates mitochondrial fission through Erk2-mediated phosphorylation of Drp1 to promote tumorigenesis (Kashatus *et al*., 2015; Serasinghe *et al*., 2015; Nagdas *et al*., 2019). Because of this, we hypothesized that KRas-mutant LUAD may be sensitive to unopposed MAPK-driven fission following inhibition of mitochondrial fusion. To inactivate mitochondrial fusion, we targeted Opa1, which is alone sufficient to prevent mitochondrial fusion. To genetically deplete Opa1, we stably expressed Cas9 along with a non-targeting sgRNA or one of two Opa1-targeting sgRNAs in KPY40 cells that we derived from a KP GEMM of LUAD and observed robust depletion of Opa1 expression (Figure 1A). To assess the effects of inhibiting Opa1 on individual cell growth and survival, we performed colony formation assays and found Opa1 depletion significantly inhibits colony formation (Figure 1B). We also confirmed this effect in human *KRAS*-mutant A549 LUAD cells (Figures 1C and 1D). To complement the genetic approach, we pharmacologically inhibited Opa1 using the Opa1 inhibitor MYLS22 (Herkenne *et al*., 2020). To validate inhibition of mitochondrial fusion, we examined mitochondrial morphology in DMSO- and MYLS22-treated mouse embryonic fibroblasts (MEFs). As expected, DMSO-treated MEFs demonstrate largely tubular mitochondrial morphology whereas MYLS22-treated cells demonstrate more-fragmented morphology (Figure 1E). To assess whether pharmacological inhibition of Opa1 inhibits KP LUAD colony formation, we treated KPY40 cells with DMSO or one of three doses of MYLS22. MYLS22-treated cells demonstrate decreased colony formation at doses of 25 uM and 50 uM (Figure 1F). Together, these data demonstrate that genetic or pharmacologic inhibition of Opa1 decreases KRas-mutant LUAD colony formation. To determine whether the effects of Opa1 inhibition are mediated by unopposed mitochondrial fission, we examined colony formation under Drp1 depletion, Opa1 depletion, or both in KPY40 cells. To generate these conditions, we first stably introduced Cas9 with a non-targeting sgRNA or one of two Drp1-targeting sgRNAs. In each of these three sets of cells, we then stably introduced a non-targeting sgRNA or an Opa1-targeting sgRNA (Figure 1G). Notably, inactivation of mitochondrial fission by Drp1 knockout does not affect colony formation in these cells (Figures 1H and 1I). This was surprising given we have previously found that Drp1 depletion inhibits KP pancreatic ductal adenocarcinoma (PDAC) growth. Opa1 depletion alone in this double-CRISPR system recapitulates the inhibition of colony formation we observed in the single-CRISPR system. Inactivation of mitochondrial fission completely rescues the effects of Opa1 knockout on colony formation. These results suggest that the effects of Opa1 knockout on *in vitro* colony formation of KP LUAD cells are dependent on mitochondrial fission and suggest a functional link between Drp1 and Opa1.

**Figure 1:**
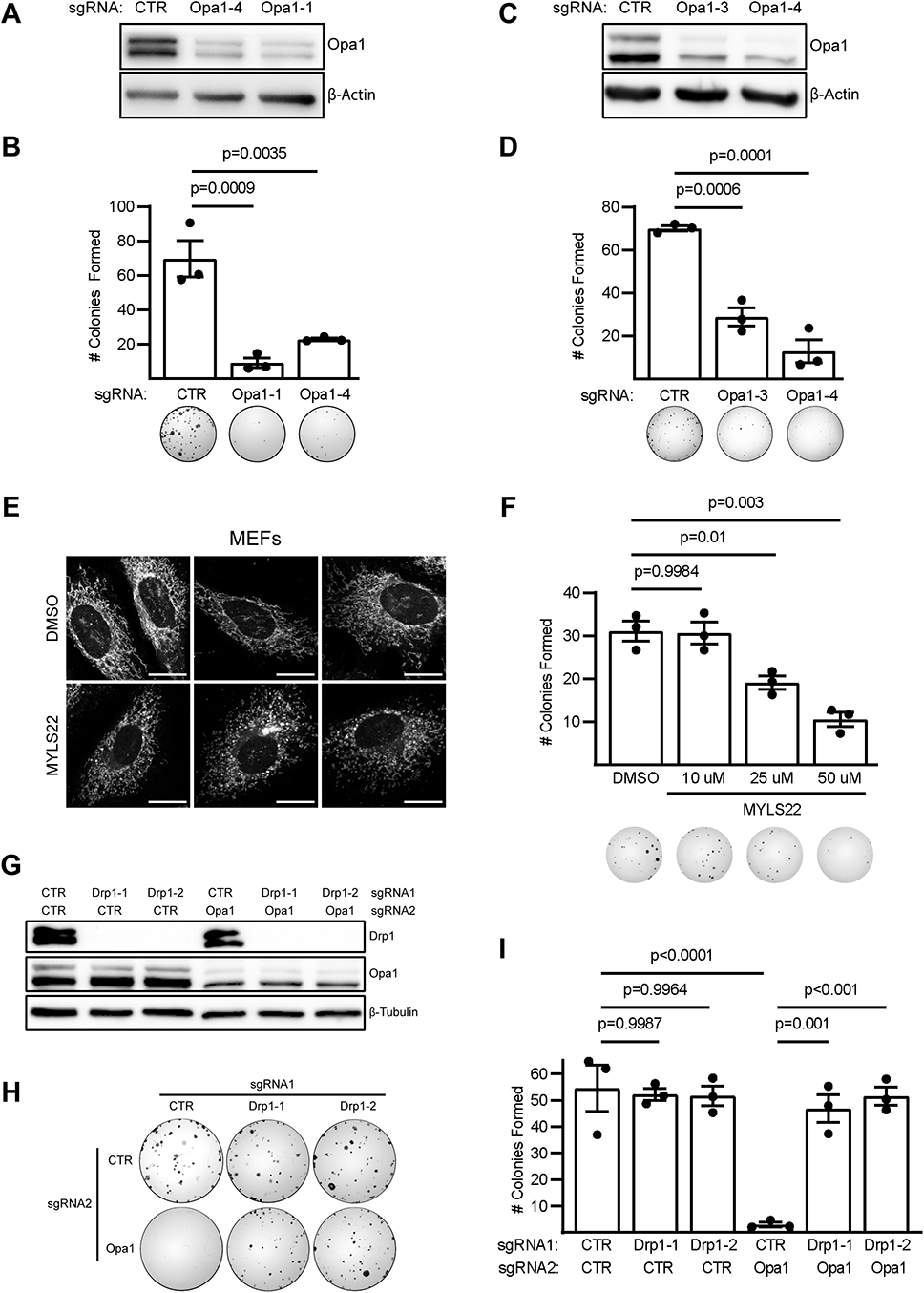
Opa1 inhibition prevents KRas*-*mutant LUAD colony formation in a Drp1-dependent manner: A. Immunoblot confirmation of CRISPR-mediated Opa1 depletion in KPY40 mouse LUAD cells. B. Representative wells from colony formation assays in indicated KPY40 CRISPR cells with quantification. n=3 independent experiments. Mean±SD. One-way ANOVA + Dunnett’s multiple comparison test. C. Immunoblot confirmation of CRISPR-mediated Opa1 depletion in A549 cells. D. Representative wells from colony formation assays in indicated A549 CRISPR cells with quantification. n=3 independent experiments. Mean±SD. One-way ANOVA + Dunnett’s multiple comparison test. E. Mitochondrial morphology in Mitotracker Red CMXRos-stained MEFs treated with DMSO or 50uM MYLS22 for 72 hours. Scale bar = 20μm. F. Representative wells from colony formation assays in MYLS22-treated KPY40 cells with quantification. n=3 independent experiments. Mean±SD. One-way ANOVA + Sidak’s multiple comparison test. G. Immunoblot confirmation of CRISPR-mediated Drp1 and/or Opa1 depletion in KPY40 cells. H. Representative wells from colony formation assays in indicated double CRISPR KPY40 cells. I. Quantification of double CRISPR KPY40 cell colony formation assays. n=3 independent experiments. Mean±SD. One-way ANOVA + Sidak’s multiple comparison test.

### Deletion of Opa1, but not Drp1, inhibits KP LUAD development *in vivo* (Figure 2)

The *in vitro* colony formation results demonstrate that inhibition of Opa1 blocks colony formation of KP LUAD cells in a Drp1-dependent manner while depletion of Drp1 alone has no effect. To test whether depletion of Drp1, Opa1, or both affects spontaneous KP LUAD development *in vivo*, we generated *KRAS^LSL-G12D/+^*; *TP53^FL/FL^* mice wildtype for *DRP1* and *OPA1* (KP), or with homozygous floxed alleles for *DRP1* (KPD), *OPA1* (KPO), or both (KPDO) (Figure 2A). We initiated tumor formation and mitochondrial dynamics gene depletion by intratracheal administration of Adenovirus-Cre (AdCre) and allowed tumors to develop for 10 weeks (Figure 2B). Contrary to what is observed in KRas-driven PDAC (Nagdas *et al*., 2019), LUAD tumor burden in KPD mice is not significantly different from that of KP (Figure 2C); however, tumor burden in KPO mice is significantly less than in KP (Figure 2C). Surprisingly, simultaneous deletion of Opa1 and Drp1 does not rescue tumor development, as tumor burden is decreased in KPDO mice compared to KP and no different from KPO. Hematoxylin- and eosin-stained lung sections demonstrate that all mice developed lung tumors with similar histological morphology (Figure 2D). Together, these data suggest that Drp1 is dispensable, but Opa1 is required, for KP LUAD development, and that deletion of Drp1 is insufficient to rescue Opa1 deletion-mediated decrease in tumor development *in vivo*.

**Figure 2:**
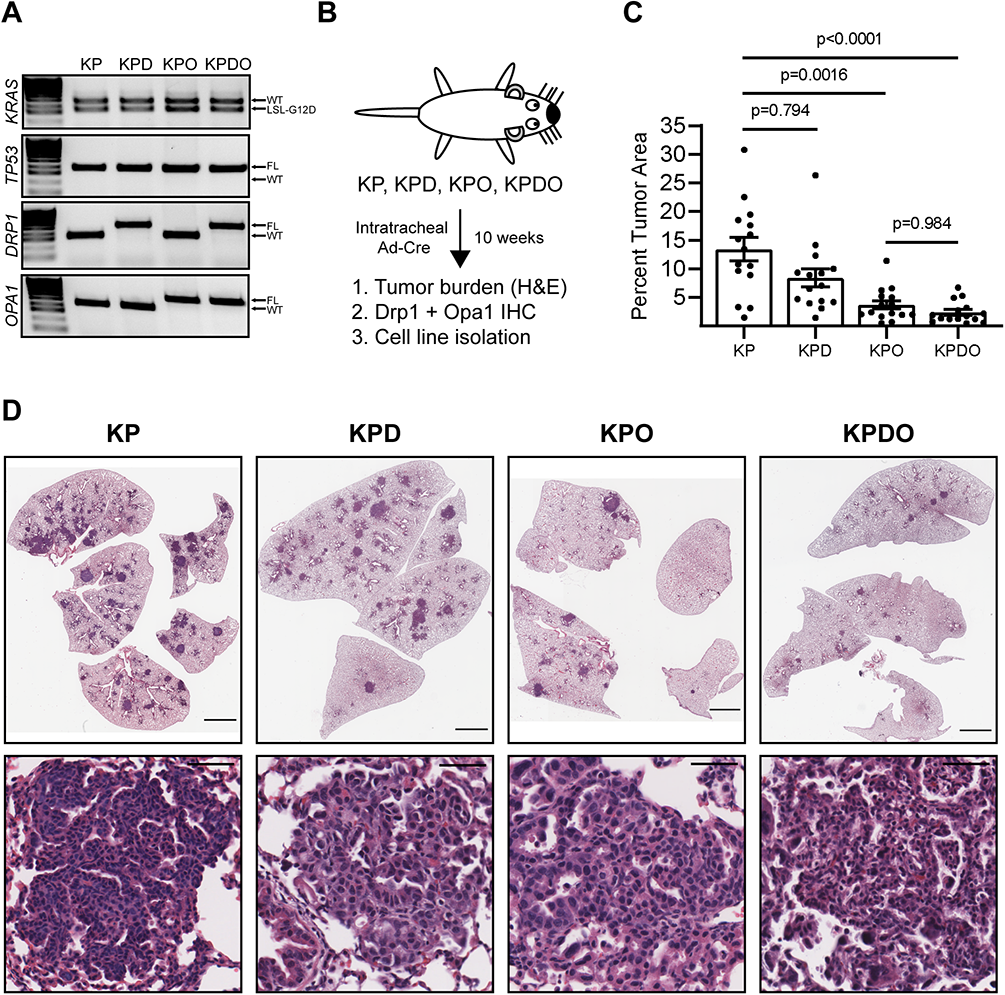
Deletion of Opa1, but not Drp1, inhibits KP LUAD development *in vivo*. A. PCR genotypes for *KRAS, TP53, DRP1,* and *OPA1* alleles in GEMM-enrolled mice B. KPDO GEMM study schematic and endpoints. C. Tumor burden in GEMM mice as percent tumor area versus total lung area. n=15 mice per genotype. Mean±SD. Kruskal-Wallis test + Dunn’s multiple comparison test. D. Representative hematoxylin- and eosin-stained lungs (top) and individual tumors (bottom) in indicated genotypes of mice. Scale bar = 2000 μm (whole lung), 50 μm (individual tumors).

### Opa1, but not Drp1, is required for KP LUAD development (Figure 3)

Recombination of floxed alleles by AdCre in the KP LUAD model has been demonstrated to be incompletely efficient (Eichner *et al*., 2019). Recombination of *KRAS^LSL-G12D^* and *TP53^FL/FL^* alleles is required for KP LUAD development in 10 weeks, but other floxed alleles may evade homozygous recombination. The degree to which developed tumors retain floxed alleles provides insight to whether their gene products are required for tumor development. For example, there is a strong selective pressure to retain Drp1 expression in a PDAC GEMM with KP and KPD genetics identical to the LUAD model used here (Nagdas *et al*., 2019). We took two approaches to determine whether individual tumors that formed in KPD, KPO, and KPDO mice completely recombined mitochondrial dynamics alleles. First, we performed immunohistochemistry (IHC) on lung sections using antibodies targeting Drp1 or Opa1. Second, we isolated independent tumor cell lines from mice to look for the presence of Drp1 and Opa1 by immunoblot and PCR. Tumors in KPD and KPDO mice demonstrate high efficiency of Drp1 deletion *in vivo* as evidenced by decreased Drp1 IHC staining intensity compared to KP mice (Figure 3A and 3B). Conversely, tumors that form in KPO and KPDO mice retain Opa1 expression similar to levels observed in KP mice (Figures 3C and 3D). Consistent with this, six of seven individually-derived KPD tumor cell lines demonstrate complete knockout of Drp1 while the seventh demonstrates decreased expression compared to KP cells (Figure 3E). Conversely, all seven independently-derived KPO tumor cell lines retain expression of Opa1, with most demonstrating decreased Opa1 expression compared to KP-derived cells (Figure 3F). Interestingly, six of nine KPDO cell lines demonstrate complete deletion of Opa1 *in vivo,* but this exclusively occurs in cell lines that also exhibit deletion of Drp1 (Figure 3G). Three additional KPDO cell lines retain expression of Opa1 with or without retained Drp1 expression. Consistent with the immunoblot analysis, PCR genotyping reveals that six of seven KPO cell lines recombined a single floxed *OPA1* allele *in vivo* and one of seven completely evaded recombination (Figure 3G), indicating that KP *OPA1^-/FL^* cells can develop tumors. Further, six of nine KPDO cell lines harbor homozygous recombined *OPA1^Δ^* alleles while the other three retain either one or both floxed *OPA1* alleles (Figure 3I). Together, these data suggest that mitochondrial fission is dispensable for KP LUAD development *in vivo*, and that Opa1 is essential in tumor cells with intact mitochondrial fission. Further, isolation of six cell lines with complete *in vivo* Opa1 deletion exclusively in the context of complete Drp1 deletion suggests that inactivation of mitochondrial fission permits cell line isolation but not *in vivo* tumor development.

**Figure 3:**
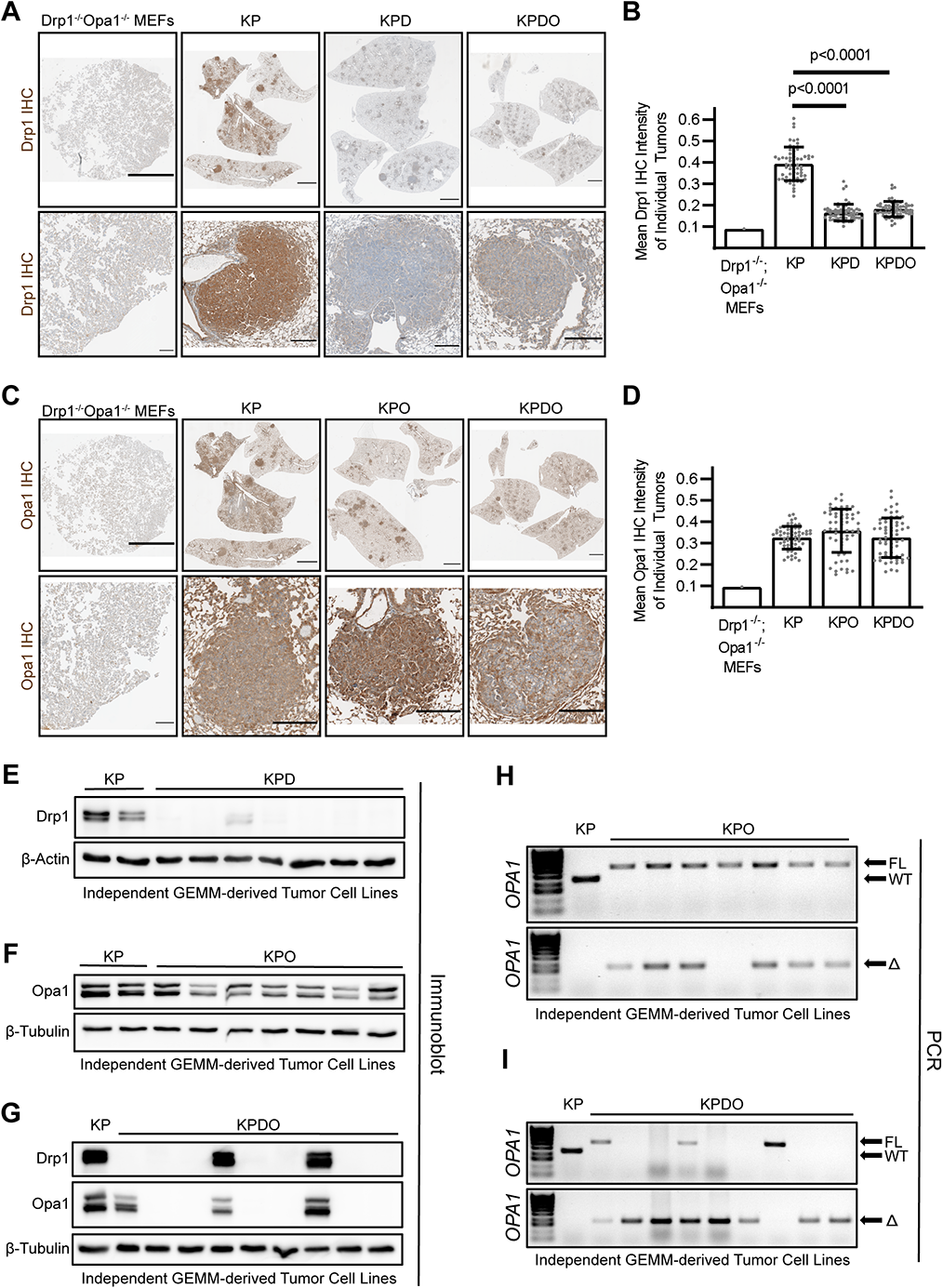
Opa1, but not Drp1, is required for KP LUAD development. A. Representative Drp1 IHC images on whole lung (top) and individual tumors (bottom) in indicated genotypes. KPDO MEFs serve as negative staining control. Scale bar = 2000 μm (whole lung), 200 μm (individual tumors). B. Quantification of mean Drp1 DAB intensity of individual tumors in indicated genotypes. n=60 tumors per genotype. Kruskal-Wallis test + Dunn’s multiple comparison test. C. Representative Opa1 IHC images on whole lung (top) and individual tumors (bottom) in indicated genotypes. KPDO MEFs serve as negative staining control. Scale bar = 2000 μm (whole lung), 200 μm (individual tumors). D. Quantification of mean Opa1 DAB intensity of individual tumors in indicated genotypes. n=60 tumors per genotype. E. Immunoblot of Drp1 expression in 2 KP and 7 KPD independently-derived tumor cell lines. F. Immunoblot of Opa1 expression in 2 KP and 7 KPO independently-derived tumor cell lines. G. Immunoblot of Drp1 and Opa1 expression in 1 KP and 9 KPDO independently-derived tumor cell lines. H. PCR genotypes for *OPA1^FL^* and *OPA1^WT^* alleles (top) and *OPA1^Δ^* (recombined, bottom) in 1 KP and 7 independently-derived KPO tumor cell lines. I. PCR genotypes for *OPA1^FL^* and *OPA1^WT^* alleles (top) and *OPA1^Δ^* (recombined, bottom) in 1 KP and 9 independently-derived KPDO tumor cell lines.

### Opa1 is required to maintain mitochondrial NAD^+^ regeneration (Figure 4)

To better understand what causes reduced tumor burden in KPO mice compared to KP, we explored the consequences of Opa1 knockout *in vitro* using a KPO tumor cell line that retained both floxed *OPA1* alleles *in vivo*. Infection of this cell line *in vitro* with AdCre, but not empty vector adenovirus (AdEmpty), deletes Opa1, whereas AdCre infection of a tumor-derived KP cell line has no effect on Opa1 expression (Figure 4A). Opa1 protein is almost completely undetectable three days after AdCre infection of KPO cells, which allows determination of the effects of acute Opa1 knockout in KP LUAD. As depletion of Opa1 has previously been reported to sensitize cells to apoptosis (Olichon *et al*., 2003; Frezza *et al*., 2006), we first assessed whether Opa1 knockout universally decreases cell viability by treating tumor cells with known apoptosis inducers cisplatin or etoposide. Surprisingly, Opa1 knockout cells demonstrate slightly higher cell viability in cisplatin and etoposide relative to DMSO compared to Opa1-expressing cells and there is no visible difference in cell accumulation (Figure S1A and S1B). Additionally, Opa1 knockout cells do not demonstrate significant induction of PARP cleavage upon cisplatin or etoposide treatment (Figure S1C). Together, these data suggest that Opa1 knockout does not sensitize KP LUAD to apoptosis. Opa1 depletion has also been reported to impair OXPHOS capacity in mouse fibroblasts (Cogliati *et al*., 2013; Del Dotto *et al*., 2017; Quintana-Cabrera *et al*., 2018; Cretin *et al*., 2021). To test whether KP LUAD cells are similarly sensitive, we performed Seahorse mitochondrial stress test assays and found that acute Opa1 knockout severely impairs oxygen consumption (OCR) in KP LUAD (Figure 4B). This effect is not due to AdCre infection itself, as there is no change in OCR in KP cells following infection (Figure 4D and 4E). ETC dysfunction following Opa1 deletion decreases ATP synthesis (Gomes, Benedetto and Scorrano, 2011; Cogliati *et al*., 2013; Quintana-Cabrera *et al*., 2018); however, recent evidence suggests that cancer cells fulfill ATP requirements through glycolysis and instead require the ETC to couple electron flux to the oxidation of the pyrimidine precursor dihydroorotate (DHO) to orotate and NADH to NAD^+^, which serves as an essential cofactor for oxidative biosynthesis (Birsoy *et al*., 2015; Sullivan *et al*., 2015; Bajzikova *et al*., 2019; Martínez-Reyes *et al*., 2020; Luengo *et al*., 2021). We thus hypothesized that Opa1 deletion-mediated ETC dysfunction impairs KP LUAD growth by inhibiting the ETC electron flux required for oxidation of electron-carrying molecules such as DHO and NADH, and that mitochondrial ATP synthesis is dispensable. To determine which ETC functions are required in this model, we treated KP LUAD cells with DMSO, oligomycin, rotenone, or CCCP and measured cell accumulation. CCCP inactivates mitochondrial ATP synthesis through dissipation of the proton gradient (Figure S1D), but leaves ETC electron flow intact (Figure S1E), whereas both rotenone and oligomycin diminish mitochondrial ATP synthesis and ETC electron flux (S1E). Both oligomycin and rotenone almost completely inhibit cellular accumulation compared to DMSO, while CCCP treatment has no effect (Figure 4F). These data suggest that ETC-mediated oxidation of electron carriers is required and that mitochondrial ATP synthesis is dispensable for KP LUAD growth. To assess whether Opa1 deletion affects NAD^+^ metabolism, we measured the ratio of NAD^+^ to NADH and found that Opa1 deletion severely reduces this ratio, indicating dysfunction of NADH oxidation (Figure 4G). Impairment of ETC electron flux causes auxotrophy for pyruvate, which supports cytoplasmic NAD^+^ regeneration through lactate dehydrogenase (LDH), and for uridine, which restores pyrimidine synthesis when DHO is unable to be oxidized (King and Attardi, 1996; Birsoy *et al*., 2015; Sullivan *et al*., 2015). To test whether the effects of Opa1 deletion on KP LUAD colony formation are mediated by inhibition of pyrimidine synthesis, NAD^+^ regeneration, or both, we treated cells with uridine alone and in combination with pyruvate and the alternative NAD^+^-regenerating LDH substrate, alphaketobutyrate (AKB). Colony formation of Opa1-retaining cells is unaffected by treatment with uridine, pyruvate, or AKB (Figure 4H), indicating that cells expressing Opa1 are able to meet metabolic demands for NAD^+^ and uridine. Notably, pyruvate and AKB supplementation increase Opa1-null colony formation, whereas uridine alone demonstrates no effect and does not increase colony formation of cells treated with pyruvate or AKB. These data suggest that Opa1 deletion impairs low-density colony formation by inhibiting NAD^+^ regeneration, but not pyrimidine synthesis. To determine whether Opa1 deletion sensitizes cells to inhibition of cytoplasmic NAD^+^ regeneration, we measured tumor cell viability in cells treated with DMSO, the lactate dehydrogenase inhibitor GNE-140 (Boudreau *et al*., 2016), or the PDK inhibitor AZD7545 that activates PDH and drives pyruvate into mitochondria and away from lactate synthesis. We found Opa1-null cells demonstrate decreased viability compared to Opa1-expressing cells when treated with GNE-140 or AZD7545 (Figure 4I). Together, these data suggest that Opa1 knockout inhibits KP LUAD by restricting mitochondrial ETC-mediated NAD^+^ regeneration.

**Figure 4:**
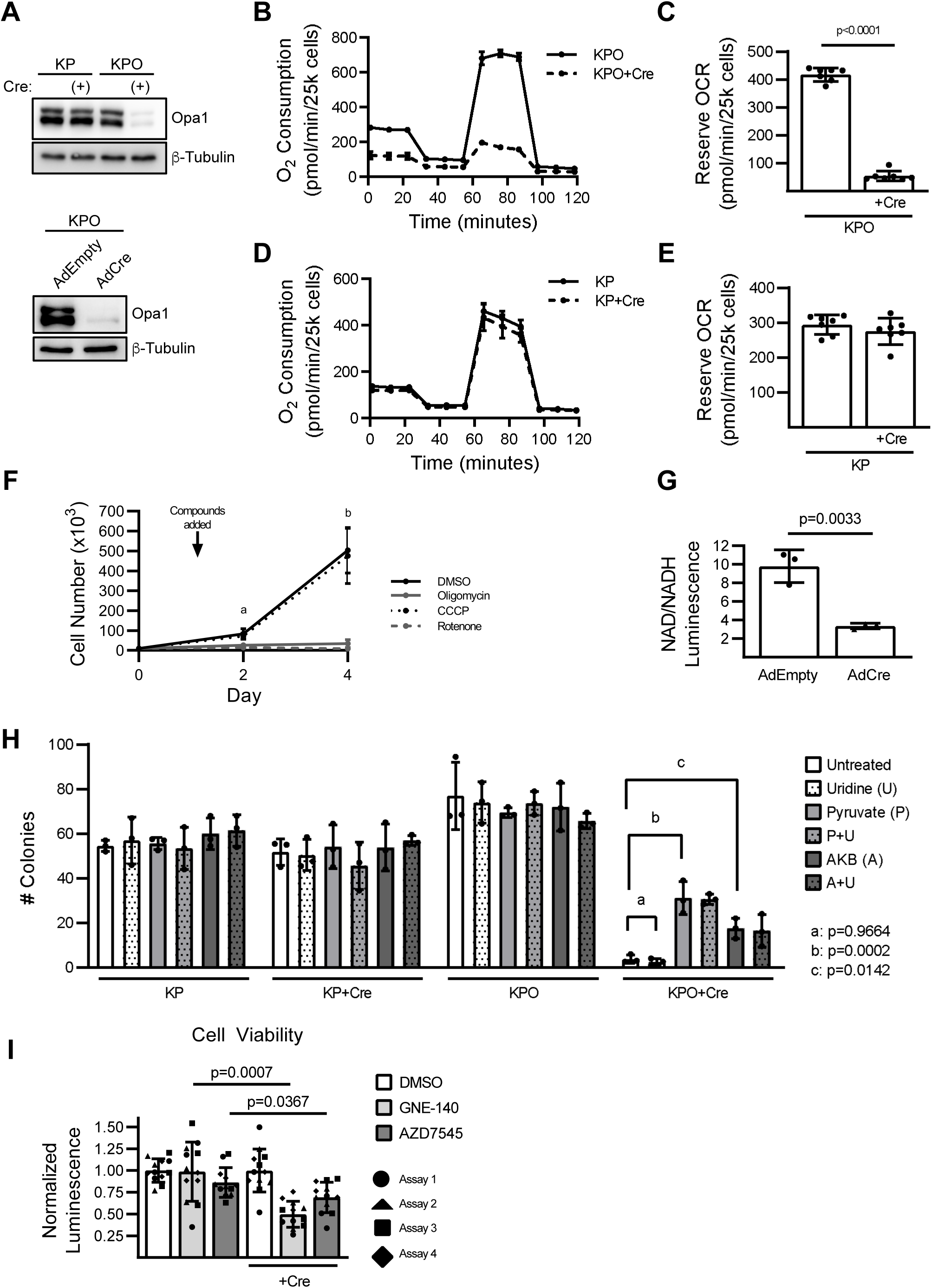
Opa1 is required to maintain mitochondrial NAD^+^ regeneration. A. Immunoblot of Opa1 expression in untreated and AdCre-infected GEMM-derived KP and KPO tumor cells (top), and AdEmpty- or AdCre-infected KPO (bottom). Lysates generated three days after adenovirus infection. B. Oxygen consumption of untreated or AdCre-infected KP cells measured by Seahorse mitochondrial stress test. n=7 wells per cell condition. Mean±SD. C. Reserve oxygen consumption of untreated or AdCre-infected KP cells measured by Seahorse mitochondrial stress test. n=7 wells per cell condition D. Oxygen consumption of untreated or AdCre-infected KPO cells measured by Seahorse mitochondrial stress test. n=7 wells per cell condition. Mean±SD. E. Reserve oxygen consumption of untreated or AdCre-infected KPO cells measured by Seahorse mitochondrial stress test. n=7 wells per cell condition. Mean±SD. Student’s T-test. F. Cell accumulation of Opa1-expressing (no Cre) KPO cells treated with DMSO, oligomycin (10 nM), CCCP (250 nM), or rotenone (250 nM). n=3 independent experiments. Mean±SD. a = DMSO vs oligomycin/rotenone (p<0.005), DMSO vs CCCP (p=0.781), One-way ANOVA + Dunnett’s multiple comparison test. b = DMSO vs oligomycin/rotenone (p<0.05), DMSO vs CCCP (p=0.989), Welch ANOVA + Dunnett’s T3 multiple comparison test. G. NAD/NADH ratios in KPO cells treated with AdEmpty or AdCre and measured by NAD/NADH-Glo. n=3 independent experiments. Mean±SD. Student’s T-test. H. Colony formation in uninfected and AdCre-infected KP and KPO cells without treatment or treated with uridine (0.1 mg/mL), pyruvate (1 mM), alphaketobutyrate (AKB, 1 mM), or a combination as indicated. n=3 independent experiments. Mean±SD. One-way ANOVA + Dunnett’s multiple comparison test. I. CellTiter-Glo measurement of cell viability in uninfected or AdCre-infected KPO cells treated with DMSO, the lactate dehydrogenase inhibitor GNE-140 (5 uM), or the PDK1 inhibitor AZD7545 (5 uM) for 48 hours. All technical replicates from independent experiments are shown. Individual wells of drug-treated (non-DMSO) cells were normalized to the mean of the DMSO-treated wells within an individual experiment. Statistical analysis was performed on the mean of normalized values from individual experiments within treatment groups such that the sample size of each treatment group was equal to the number of independent experiments. n=4 independent experiments. Mean±SD. Student’s T-test.

### Drp1 activity drives Opa1 deletion-mediated ETC dysfunction (Figure 5)

The effect of Opa1 knockout on KP LUAD colony formation is dependent on Drp1 (Figure 1H and 1I). Although Drp1 deletion does not rescue *in vivo* tumor growth following Opa1 deletion, we were able to isolate KPDO tumor cell lines with complete *in vivo* deletion of both Opa1 and Drp1 but not a single KPO cell line that deleted only Opa1 *in vivo*. This suggests that there may be an advantage to inactivation of mitochondrial fission in the context of Opa1 deletion. To test whether co-deletion of Drp1 and Opa1 in KP LUAD affects ETC function and NAD^+^ regeneration, we utilized a KPDO tumor cell line that retained wildtype levels of Drp1 and Opa1 (Figure 3G). AdCre infection of this cell line deletes Drp1 and Opa1 within three days of infection, whereas AdEmpty infection does not (Figure 5A). We performed Seahorse mitochondrial stress test assays in these cells and found that mitochondrial oxygen consumption is unaffected by simultaneous deletion of Drp1 and Opa1 (Figure 5B and 5C). This was surprising given the severe depletion of oxygen consumption capacity we observe from Opa1 deletion alone (Figure 4D and 4E). We then tested whether co-deletion of Drp1 and Opa1 affects the NAD^+^/NADH ratio and again observed no effect (Figure 5D). These data confirm that the effects of Opa1 knockout on KP LUAD ETC function and NAD^+^ regeneration absolutely require Drp1. To determine whether the GTPase activity of Drp1 is required to mediate the effects of Opa1 deletion, we expressed luciferase, wildtype mouse Drp1 (mDrp1 WT), or the GTPase-inactive mDrp1 K38A in another KPDO tumor cell line that deleted Drp1, but not Opa1, *in vivo* (Figure 5E). AdCre infection of these cells deletes the remaining Opa1, although we observed selection for retaining Opa1 expression in mDrp1 WT-, but not mDrp1 K38A-expressing cells following AdCre infection. We performed colony formation assays in these cells and found that mDrp1 WT-expressing tumor cells form fewer colonies than control luciferase-expressing cells (Figure 5F). Additionally, tumor cells expressing mDrp1 K38A form significantly more colonies than cells expressing mDrp1 WT, indicating that the Drp1 GTPase function required for mitochondrial fission is also required to mediate the effects of Opa1 deletion on colony formation. These data indicate that Opa1 is dispensable for KP LUAD ETC function and NAD^+^ regeneration when Drp1-driven fission is inactive. Of note, these experiments tested cells under acute Opa1 and Drp1 deletion but do not provide insight into the effects of long-term inhibition of fusion-fission dynamics. As such, we next used three independent KPDO cell lines that completely knocked out Opa1 and Drp1 expression *in vivo* to test whether chronic deletion of Drp1 and Opa1 affects ETC function compared to three independent KP tumor cell lines (Figure 5G). We found that chronic deletion of Drp1 and Opa1 significantly reduces oxygen consumption (Figure 5H and 5I). This suggests that although Drp1 deletion rescues mitochondrial function from acute Opa1 deletion, chronic deletion leads to decreased mitochondrial function and may explain why tumor burden in KPDO GEMM mice was no different than that of KPO mice. Together, these data demonstrate that Drp1 activity is required to mediate ETC dysfunction caused by acute Opa1 knockout in KP LUAD, but that long-term loss of dynamic fusion-fission cycling leads to decreased ETC function in the context of chronic Opa1 and Drp1 deletion.

**Figure 5:**
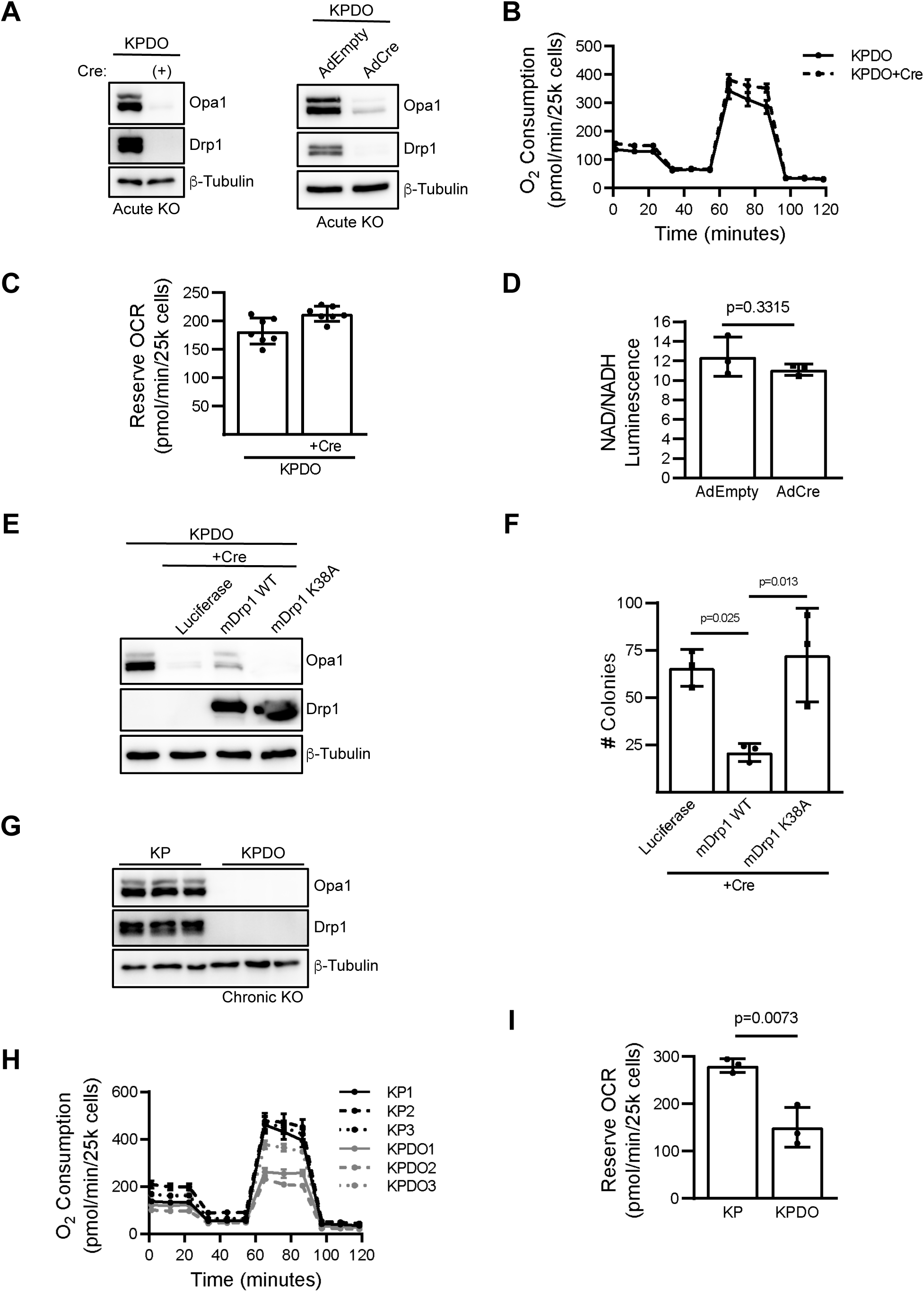
Drp1 activity drives Opa1 deletion-mediated ETC dysfunction. A. Immunoblot of Drp1 and Opa1 expression in untreated and AdCre-infected GEMM-derived KPDO tumor cells (left), and AdEmpty- or AdCre-infected KPDO (right). Lysates generated three days after adenovirus infection. B. Oxygen consumption of untreated or AdCre-infected KPDO cells measured by Seahorse mitochondrial stress test. n=7 wells per cell condition. Mean±SD. C. Reserve oxygen consumption of untreated or AdCre-infected KP cells measured by Seahorse mitochondrial stress test. n=7 wells per cell condition D. NAD/NADH ratios in KPDO cells treated with AdEmpty or AdCre and measured by NAD/NADH-Glo. n=3 independent experiments. Mean±SD. Student’s T-test. E. Immunoblot of Drp1 and Opa1 expression in a GEMM-derived KPDO tumor cell line that knocked out Drp1 in vivo but retained Opa1 expression. These cells were transduced to stably express luciferase, wildtype mouse Drp1 (mDrp1 WT), or GTPase-inactive mDrp1 K38A and were AdCre infected to knockout Opa1. F. Colony formation of AdCre-infected cells in (E). n=3 independent experiments. Mean±SD. One-way ANOVA + Sidak’s multiple comparison test. G. Immunoblot of Drp1 and Opa1 expression in untreated GEMM-derived KP and KPDO cell lines. KPDO cells deleted Drp1 and Opa1 *in vivo.* n=3 independently-derived cell lines per genotype. H. Oxygen consumption of KP and KPDO cells measured by Seahorse mitochondrial stress test. n=3 independent cell lines per genotype. Mean±SD. I. Mean reserve oxygen consumption of KP and KPDO cell lines measured by Seahorse mitochondrial stress test. n=7 wells for each of 3 independent cell lines per genotype. Mean±SD. Student’s T-test.

### Drp1 mediates ETC disassembly and dysmorphic cristae following Opa1 knockout (Figure 6)

Our results suggest that Drp1 activity drives ETC dysfunction in the context of acute Opa1 deletion. Since the ETC is embedded in the mitochondrial inner membrane and its function is promoted by proper cristae morphology (Cogliati *et al*., 2013), we hypothesized that the interaction between Opa1 and Drp1 that drives ETC function affects cristae homeostasis as opposed to mitochondrial outer membrane morphology. Although Opa1 is a key regulator of cristae remodeling (Olichon *et al*., 2003; Frezza *et al*., 2006; Anand *et al*., 2014; Del Dotto *et al*., 2017), its exact function in this context is unclear. The role of Drp1 in cristae remodeling is also unclear, but there is evidence that fission promotes cristae disorganization that increases cytochrome C release from mitochondria during intrinsic apoptosis (Otera *et al*., 2016). We therefore hypothesized that Drp1 disrupts cristae organization at steady state, not only during apoptosis, and that Opa1 opposes Drp1 by reorganizing cristae following fission events. To assess how Opa1 deletion affects ETC function and cristae homeostasis in the presence and absence of Drp1, we again used KPO and KPDO tumor cell lines that retain floxed *OPA1* or *OPA1* and *DRP1* alleles. *In vitro* AdCre infection deletes remaining floxed alleles in these cells, whereas AdEmpty does not (Figure 6A). Since we observed a significant decrease in the NAD^+^/NADH ratio in Opa1-deleted cells that retain Drp1, we assessed the assembly of ETC complexes and the activity of ETC complex I that oxidizes NADH to NAD^+^. To determine whether Opa1 deletion inhibits complex I assembly in a Drp1-dependent manner, we performed clear native PAGE (cnPAGE) on mitochondrial isolates from AdEmpty- and AdCre-infected KPO and KPDO tumor cells and visualized ETC complexes with Coomassie stain. Opa1 deletion substantially decreases levels of assembled complexes I, III, and V, whereas co-deletion of Opa1 and Drp1 almost completely rescues this effect (Figure 6B). We next assessed *in vitro* complex I activity in these samples using cnPAGE followed by a complex I in-gel activity (IGA) assay and found that Opa1 deletion decreases complex I activity and that co-deletion with Drp1 rescues this effect (Figure 6C and 6D). Equal loading of mitochondrial isolates for ETC assembly and complex I IGA assays was confirmed by immunoblot for mitochondrial proteins SDHA and VDAC (Figure 6E). These data suggest that Drp1-expressing cells require Opa1 to maintain ETC complex I assembly and function, but that cells lacking Drp1 are able to assemble functional ETC complexes in the absence of Opa1. Since loss of Opa1 causes depletion of mitochondrial DNA and disrupts cristae morphology (Olichon *et al*., 2003; Elachouri *et al*., 2011; Del Dotto *et al*., 2017), we reasoned that either decreased expression of mitochondrial DNA-encoded ETC subunits or disrupted cristae morphology could explain the loss of ETC complex assembly and function following acute deletion of Opa1 in KP tumor cells. To address this, we analyzed cristae structure and measured the abundance of mitochondrial DNA in AdEmpty- or AdCre-treated KPO and KPDO tumor cells. Transmission electron microscopy (TEM) analysis of cristae morphology in these cells revealed that lamellar cristae, in which the cristae membrane contacts the inner boundary membrane (IBM), are abundant in the mitochondria of Opa1-expressing cells. Opa1 knockout decreases the proportion of mitochondria with lamellar cristae and increases the proportion of mitochondria with tubular or no discernable cristae but this phenotype is not observed following knockout of both Opa1 and Drp1 (Figure 6F, 6G, 6H and S2). These data indicate that mitochondrial fission activity drives loss of cristae structure following Opa1 deletion. We next assessed the impact of Opa1 deletion on the relative abundance of mitochondrial DNA versus nuclear DNA in these cells by measuring the levels of mitochondrial-encoded genes *ND1* and *16S* and the nuclear-encoded gene *HK2* by quantitative PCR. Acute Opa1 deletion reduces abundance of mtDNA to about two-thirds that of Opa1-expressing cells, whereas Opa1 deletion has no effect on mtDNA abundance when Drp1 is also deleted (Figure 6I and 6J). Although the decrease in mtDNA abundance from acute Opa1 deletion in Drp1-expressing cells is clear, the impact of this decrease on protein expression of mtDNA-encoded ETC subunits is uncertain, especially under the acute deletion timescales of these experiments. These data suggest that at steady state, mitochondrial fission drives dysmorphic cristae structure, mtDNA loss, and decreased ETC function if left unopposed by Opa1.

**Figure 6:**
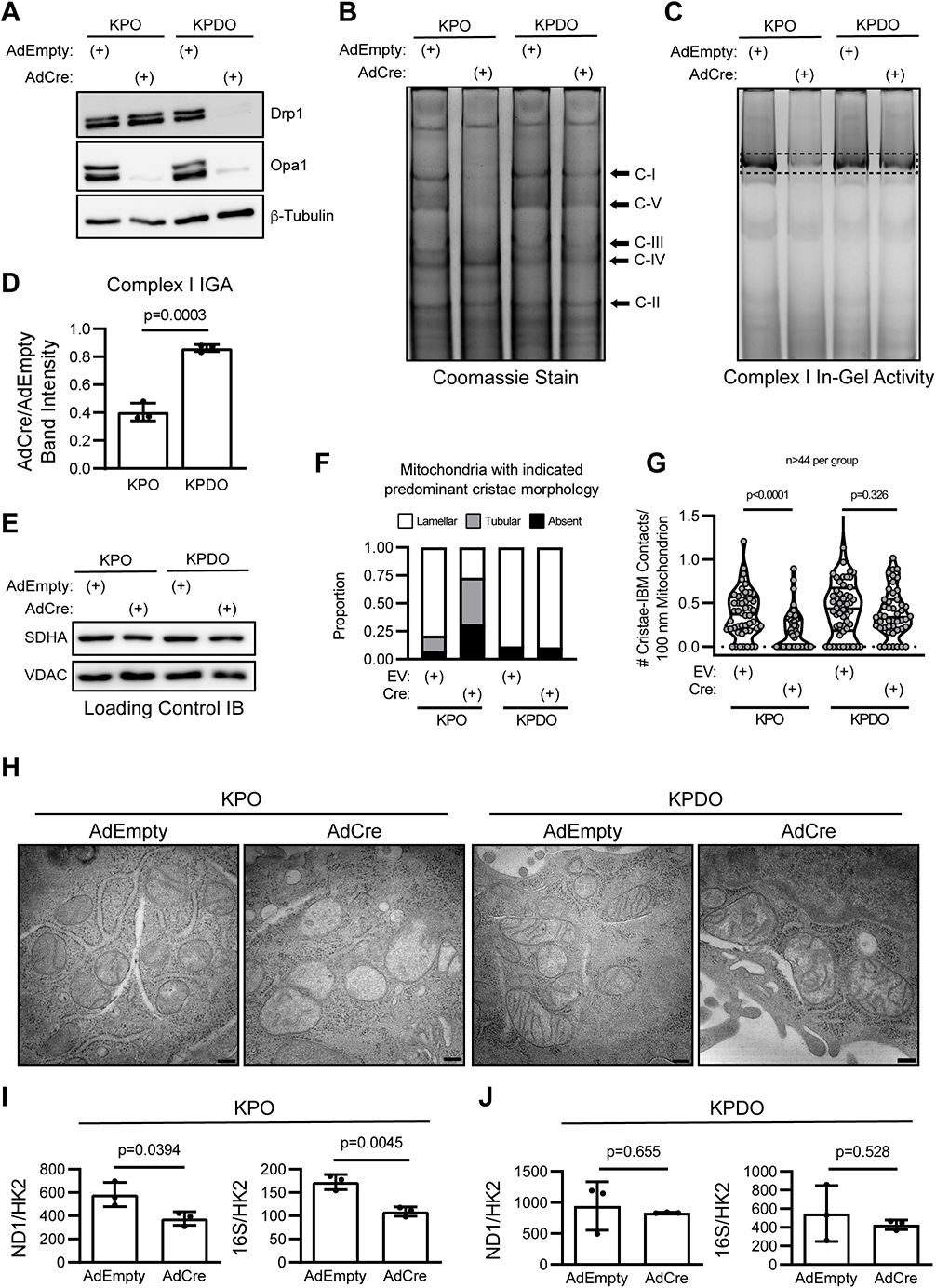
Drp1 mediates ETC disassembly and dysmorphic cristae following Opa1 knockout. A. Immunoblot of Drp1 and Opa1 expression in AdEmpty- and AdCre-infected GEMM-derived KPO and KPDO tumor cells. Lysates were generated three days after *in vitro* adenovirus infection. B. Coomassie-stained clear native PAGE of mitochondrial isolates from AdEmpty- and AdCre-infected KPO and KPDO cells (representative of 3 independent mitochondrial fractionation experiments). C-I = ETC complex I, C-V = complex V, C-III = complex III, C-IV = complex IV, C-II = complex II. C. Clear native PAGE + complex I in-gel activity (IGA) assay of mitochondrial isolates from AdEmpty- and AdCre-infected KPO and KPDO cells (representative of 3 independent mitochondrial fractionation experiments). D. Quantification of complex I IGA intensity assays. n=3 independent experiments. Mean±SD. Student’s T-test. E. Immunoblot of SDHA and VDAC for confirmation of equal mitochondrial protein loading in native PAGE experiments (representative of 3 independent mitochondrial fractionation experiments). F. Transmission electron microscopy (TEM) quantitation of dominant cristae morphology (lamellar, tubular, or absent) in individual mitochondria from AdEmpty (EV) or AdCre-treated KPO and KPDO tumor cells. G. TEM quantitation of number of cristae contacting the inner boundary membrane (IBM) per 100 nm per mitochondrion in AdEmpty or AdCre-treated KPO and KPDO tumor cells. n>44 mitochondria analyzed per cell type. Violin plot shows quartiles and median. Mann-Whitney test. H. Representative TEM images of mitochondria and cristae morphology in AdEmpty- or AdCre-infected KPO and KPDO tumor cells. Magnification = 30k. Scale bar. = 200 nm. See supplemental figure 2 for additional images. I. Quantitative PCR measuring the abundance of mitochondrial-encoded genes *ND1* (left) and *16S* (right) versus the nuclear gene *HK2* in AdEmpty- or AdCre-treated KPO tumor cells. n=3 independent experiments. Mean±SD. Student’s T-test. J. Quantitative PCR measuring the abundance of mitochondrial-encoded genes *ND1* (left) and *16S* (right) versus the nuclear gene *HK2* in AdEmpty- or AdCre-treated KPDO tumor cells. n=3 independent experiments. Mean±SD. Student’s T-test.

## Discussion

This work demonstrates that Opa1 is required *in vitro* and *in vivo* for KRas-mutant lung adenocarcinoma growth and development by promoting the ETC-mediated NAD^+^ regeneration necessary to maintain oxidative biosynthesis. We find the cell growth and metabolic phenotypes that arise from Opa1 knockout are completely reversible *in vitro* by simultaneous deletion of Drp1 or inactivation of its GTPase catalytic domain. This indicates that mitochondrial fission that is unopposed by Opa1 action is catastrophic to mitochondrial function and ultimately cell growth and survival. In stark contrast, deletion of Drp1 alone demonstrates no effect on KP LUAD tumor cell growth *in vitro* or on tumor development *in vivo*. This is surprising given the requirement for Drp1 in a PDAC model with identical KP genetics (Nagdas *et al*., 2019), and in *BRAF-*mutant melanoma, another MAPK-driven tumor system (Serasinghe *et al*., 2015). This suggests that the impacts of disrupting mitochondrial dynamics are specific to the tissues from which tumors arise, which will be critical to evaluating the potential efficacy of inhibiting mitochondrial dynamics machinery as future therapy for human disease.

Results from the KP GEMM indicate that Opa1 is required for *in vivo* tumor development, as KPO mice demonstrate decreased tumor burden compared to KP and our inability to establish tumor cell lines with homozygous *in vivo* Opa1 knockout; however, six out of seven KPO tumor cell lines demonstrated *OPA1^FL/Δ^* genotypes, indicating that tumors can develop from initiating cells with only one intact *OPA1* allele. In contrast to the *in vitro* experiments, simultaneous knockout of Drp1 and Opa1 is unable to rescue *in vivo* tumor burden, but permits isolation of KPDO cell lines with complete recombination of all floxed *DRP1* and *OPA1* alleles. Importantly, ETC function in cells that knocked out Drp1 and Opa1 *in vivo* is diminished compared to KP tumor cell lines, which may explain why tumor burden is not rescued *in vivo.* These data suggest that cells that with Drp1 and Opa1 deletion *in vivo* retain enough OXPHOS capacity to survive the duration of the GEMM and tissue harvest, and then proliferate *in vitro* under conditions of high nutrient availability. This also suggests that long term simultaneous inhibition of Drp1 and Opa1 is deleterious to mitochondrial function through mechanisms not examined in this work, but potentially due to decreased turnover of damaged mitochondria, decreased mitochondrial content mixing, mitochondrial DNA loss, or impaired lipid transport and synthesis, as has been recently reported under these conditions in yeast (Kojima *et al*., 2019).

Opa1 is likely required by all tumor types and tissues that require efficient ETC function, but the specific ETC function required may be different in different contexts. For instance, skeletal muscle likely depends on Opa1 for efficient ETC-mediated synthesis of ATP (Lodi *et al*., 2011), whereas most tumors require the ETC to regenerate NAD^+^ to fuel oxidative biosynthesis of nucleotides and amino acids while generating ATP through glycolysis. Recent work has found that mitochondrial ATP synthesis comes at the expense of rapid NAD^+^ regeneration, as generation of the proton gradient required by mitochondrial ATP synthase slows ETC electron flux and thus NADH oxidation (Luengo *et al*., 2021). It would therefore be interesting to assess whether tumor cells that require ETC-mediated NAD^+^ regeneration, but not mitochondrial ATP synthesis, regulate Drp1 and Opa1 in a manner that promotes NADH oxidation by complex I and increases ETC electron flux, potentially through a mitochondrial fission-mediated increase in proton leakage to counteract the slowing of ETC electron flux.

Although Opa1 function may be important for many or all cell types, fine-tuning its function through pharmacological inhibition, as opposed to complete knockout, may offer therapeutic value for highly proliferative tumors with substantial ETC-mediated NAD^+^ regeneration requirements combined with highly active mitochondrial fission, such as those with activating mutations in *KRAS* or other MAPK activators. Given KP LUAD can develop with heterozygous deletion of Opa1 and decreased Opa1 protein expression (Figure 3F and 3H), we suspect that the degree of Opa1 inhibition required for therapeutic efficacy would have to exceed 50%. It is possible that ETC function in fission-stimulated tumors would be more adversely affected by Opa1 inhibition than tissues without high ETC function requirements and without fission-activating mutations, especially in the context of acute treatment. In support of this, mouse hepatocytes tolerate complete Opa1 depletion *in vivo* and Opa1 silencing reverses liver steatosis characterized by formation of megamitochondria through mitochondrial fusion (Yamada *et al*., 2022). We found that KP LUAD is sensitive to treatment with the first-in-class Opa1 inhibitor MYLS22; it would be interesting to determine whether Opa1 inhibition may offer a therapeutic window in which ETC dysfunction is more severe and more deleterious to tumor cells than normal tissue.

Multiple groups have demonstrated that Drp1 depletion induces proteolytic cleavage of Opa1 through the action of the inner mitochondrial membrane protease Oma1 (Möpert *et al*., 2009; Murata *et al*., 2020). These studies establish a regulatory nexus between mitochondrial fission and Opa1 and provide evidence of communication between these two key mitochondrial shaping proteins. However, whether and how Opa1 actively senses fission activity or actively regulates Drp1 activity is not known. It will be interesting to explore how the activities of these proteins are sensed and whether they lead to reciprocal changes in activity through posttranslational modification, localization or expression.

Opa1, the mitochondrial contact site and cristae organizing system (MICOS) complex, and ATP synthase dimers have each been implicated in the regulation of cristae morphology (Paumard, 2002; Olichon *et al*., 2003; Strauss *et al*., 2008; Harner *et al*., 2011; Hoppins *et al*., 2011; Bohnert *et al*., 2015; Varanita *et al*., 2015; Blum *et al*., 2019); however, the exact functions of, and interactions between systems that shape cristae remain unclear. Our work suggests that mitochondrial fission events disrupt cristae structure and that these disruptions must be repaired to maintain ETC function. The sequence of action and exact localization of Opa1, MICOS complex proteins, and ATP synthase dimers in establishing and maintaining cristae morphology remain unclear, though recent work suggests that Opa1 is epistatic to the core MICOS complex protein, MIC60, in promoting cristae structure (Glytsou *et al*., 2016). MIC60 densely populates cristae junctions, whereas Opa1 localizes to cristae membranes and the inner boundary membrane (Barrera *et al*., 2016). These observations suggest that Opa1 is required to initially form new cristae by directly folding the inner mitochondrial membrane while the MICOS complex maintains cristae structure by securing cristae junctions to the outer mitochondrial membrane and inhibiting Opa1 from fusing opposing leaflets of the inner mitochondrial membrane at cristae junctions. This model is consistent with a requirement for Opa1 only when Drp1-dependent fission activity disrupts cristae structure. The model also predicts that the MICOS complex is required to anchor cristae even in the absence of Drp1. Future work will delineate how fission and the MICOS complex interact to affect cristae structure. Additionally, the role of the long and short Opa1 isoforms in the formation and maintenance of cristae structure remains unclear, as some reports demonstrate that cells with the short form alone are able to maintain normal cristae morphology (Del Dotto *et al*., 2017) and others demonstrate severely disrupted cristae structure (Wunderlich *et al*., 2008). Whether Ras-induced fission promotes changes in Opa1 processing and whether expression of different Opa1 isoforms is able to promote ETC function in Opa1 null LUAD cells could provide important insights about their respective roles.

Notably, we find that acute Opa1 deletion impacts both cristae morphology and mtDNA content in a Drp1-dependent manner. We cannot rule out the possibility that Drp1 disrupts cristae folding indirectly through direct loss of mtDNA, which encodes OXPHOS complex subunits including ATP synthase. Decreased protein expression of mitochondrially-encoded ATP synthase subunits would decrease assembly and dimer formation, which could lead to a loss of membrane folding. In this alternative model, Opa1 activity would preserve cristae structure in the presence of high fission activity by preventing the fission-induced loss of mtDNA.

Collectively, this work establishes the dependency of LUAD cells on mitochondrial ETC-mediated NAD^+^ regeneration and elucidates how Opa1 opposes Drp1-mediated mitochondrial fission to promote ETC assembly and function through inner membrane restructuring. This work supports a need for future research that will inform the approach and efficacy of inhibiting mitochondrial dynamics in specific tumor types and the development of more potent and specific inhibitors of dynamics effectors.

## Acknowledgments

We thank Sheri VanHoose and the UVA Research Histology Core for help with GEMM lung FFPE preparation and sectioning; Pat Pramoonjago and the UVA Biorepository and Tissue Research Facility for help with the GEMM tumor IHC; Natalia Dworak for help with the TEM; Stacy Criswell and Adrian Halme for help with confocal microscopy.

## Author Contributions

Conceptualization: D.T.S, M.W.M., D.F.K, Methodology: D.T.S and D.F.K, Formal Analysis: D.T.S, J.A.K., Investigation: D.T.S., K.K., N.C., Resources: J.A.K., K.P, H.S., D.F.K., Writing – Original Draft: D.T.S and D.F.K., Writing – Review & Editing: D.T.S., D.F.K., M.W.M., J.A.K., H.S., Visualization: D.T.S, Funding Acquisition: D.F.K

## Declaration of Interests

The authors declare no competing interests

## Methods

### KPDO genetically-engineered mouse model (GEMM) of lung adenocarcinoma

This is a fixed endpoint study assessing tumor burden and retention of mitochondrial dynamics gene expression in KP, KPD, KPO, and KPDO mice ten weeks after intratracheal adenovirus-Cre administration. The experimental unit for analysis of tumor burden is the mouse and the sample size is 15 mice per group. The experimental unit for retention of Drp1 and/or Opa1 expression is the individual tumor nodule (IHC, n=60 per group) or individual GEMM-derived tumor cell line (immunoblot and PCR, sample size variable based on number of independent tumor cell lines generated per mouse genotype). The only inclusion criterion was desired genotype of mice as determined by PCR. All mice that satisfied genotype requirements were enrolled until the sample size within a given group was satisfied. Four KPDO mice past the desired sample size (chronologically the last four) were mistakenly enrolled and infected with adenovirus-Cre. These mice were excluded from tissue harvest for tumor burden analysis to maintain equal group sample sizes, but were used to generate GEMM-derived tumor cell lines in the same manner as mice used as intended for tumor burden analysis. No randomization was employed as mice were generated and enrolled based solely on genotype. Mice from both sexes were used in each of the four groups as generated, and the number of each sex is as follows (male, female): KP (9,6), KPD (6,9), KPO (13,2), KPDO (7,8). KP (*LSL-KRAS^G12D/+^; TP53^FL/FL^*) mice were supplied by Dr. Kwon Park. *OPA1^FL/FL^* (O) mice (Zhang *et al*., 2011) were supplied by Dr. Hiromi Sesaki. KP mice were mated with *OPA1^FL/FL^* and *DRP1^FL/FL^* (D) (Wakabayashi *et al*., 2009) to generate heterozygous KPO and KPD genotypes. KPD and KPO were mated to generate heterozygous KPDO mice. Breeders for all conditions were descendants from KPDO heterozygotes to maintain recent shared ancestry. KPD, KPO, and KPDO mice were generated by breeding parents heterozygous for floxed *DRP1* and/or *OPA1* alleles such that KP controls were generated in the same litters. Male and female mice of desired genotypes were generated from multiple breeder pairs per condition. Primer sequences used for mouse genotyping can be found in table S1. Intratracheal adenovirus-Cre infections were performed as previously described. (DuPage, Dooley and Jacks, 2009). Briefly, at ten weeks of age, enrollee mice were anesthetized with 250 mg/kg Avertin and intratracheally infected with 2.5×10^7^ PFU Adeno-Cre (Baylor Viral Vector Core). Mice were weighed and monitored twice per week for symptoms of disease including weight loss, hyperpnea, and piloerection for ten weeks. At ten weeks post-infection, mice were sacrificed by Avertin anesthetic overdose and exsanguination and lungs were harvested for fixation and/or tumor cell line isolation. At sacrifice, lungs were perfused intratracheally with ice cold 10% neutral buffered formalin. Mice from which tumor cell lines were derived had the right middle lobe ligated, removed, minced, and seeded into culture medium before perfusion. Perfused lungs were dissociated from the thorax and fixed in tubes containing formalin at 4C. Fixed lungs were paraffin embedded and sectioned to 5 um thick by the University of Virginia Research Histology Core. Hematoxylin and eosin staining and immunohistochemistry were performed by the University of Virginia Biorepository and Tissue Research Facility. All animal studies were performed in accordance with the University of Virginia Institutional Animal Care and Use Committee. The following parameters were assessed: tumor burden per mouse (measured by tumor surface area versus total lung area on H/E-stained FFPE sections), individual tumor Opa1/Drp1 expression by IHC on FFPE sections, and Opa1/Drp1 expression in GEMM-derived tumor cell lines by immunoblot and PCR. GEMM tumor burden analysis: hematoxylin- and eosin-stained (H&E) lung sections were scanned at high resolution on Aperio ScanScope to generate digital images. Tumor burden was calculated using QuPath software (Bankhead *et al*., 2017) as percent tumor area (tumor surface area/total lung area). Total lung area, excluding exterior connective tissue, and tumor area were traced using the wand tool. Total lung area per sample was calculated as the sum of the area of each individual lobe per sample. Total tumor area per sample was calculated as the sum of the area of individual tumors present per sample. Representative whole-lung H&E images were pixel-downsized 10x from original and imported into FIJI. GEMM IHC analysis: IHC lung sections were scanned at high resolution on Aperio ScanScope to generate digital images. IHC DAB intensity was analyzed using QuPath(Bankhead *et al*., 2017) on 20 individual tumors per slide on 3 Drp1- or Opa1-stained slides per genotype assessed (60 total per genotype). All lobes present on slides were sampled. Each tumor had mean DAB intensity measured. Drp1^-/-^;Opa1^-/-^ MEFs were analyzed by removing background first to exclude space between cells and then had mean DAB intensity measured. Statistical analysis of tumor burden and Drp1 IHC intensity was performed by Kruskal-Wallis one-way analysis of variance followed by Dunn’s multiple comparisons test using Prism v7 software. Descriptive statistics of tumor burden results by genotype (mean±SD): KP (13.43±7.92), KPD (8.37±6.13), KPO (3.68±2.86), KPDO (2.36±1.89). DS, KP, and KK performed anesthesia and intratracheal infection for all mice. DS monitored mice throughout study and performed sacrifice, tissue harvest, and data analysis.

### Cell Culture

Human A549 and mouse LUAD cells were cultured in high-glucose DMEM (Life Technologies #11965-092) supplemented with 10% FBS and 1% penicillin/streptomycin (“full DMEM”) at 37C in a 5% CO2 humidified incubator. For indicated uridine-containing media experiments, full DMEM was supplemented with 0.1 mg/mL uridine (Fisher Scientific).

### Mouse Embryonic Fibroblast Cell Line Generation

MEFs were prepared as previously described (Xu, 2005). Briefly, *TP53^FL/FL^* (P) mice or *KRAS^LSL-G12D/+^*^;^ *TP53^FL/FL^;DRP1^FL/FL^;OPA1^FL/FL^* (KPDO) mice were mated. At 13.5-14.5 days post-fertilization, pregnant females were euthanized by CO_2_ asphyxiation and cervical dislocation and uterine horns were isolated. Individual embryos were placed in separate dishes and had head and red organs removed with a scalpel and forceps. The remaining tissue was minced and enzymatically digested with trypsin at 37C for 10 minutes and placed into cell culture in full DMEM. Cell genotype was verified by PCR. Cells were immortalized by *in vitro* Adeno-Cre infection that facilitated homozygous recombination and deletion of the *TP53* alleles. P MEFs were used for Mitotracker Red staining (Figure 1). KPDO MEFs were used to validate and optimize Drp1 and Opa1 IHC staining for use on GEMM lung sections.

### Colony Formation Assay

100 cells were seeded per well in triplicate in 12-well plates and grown for 1 week in full DMEM. After one week, cells were rinsed once in PBS and fixed in 10% formalin for 20 minutes at room temperature. Formalin was aspirated and cells were stained with 0.5 mL per well of 0.5% crystal violet solution (0.5g crystal violet, 20 mL methanol, 80 mL H20) for 10 minutes at room temperature. Crystal violet solution was removed and cells were rinsed in H_2_0 twice for 10 minutes on a shaker before drying and imaging. Colony number was quantified using FIJI.

### CellTiterGlo (CTG) Cell Viability Assay

500 cells were seeded in 50 uL full DMEM in white-walled 96-well plates in technical duplicates or triplicates per independent experiment. The next day, media was changed to 50uL fresh DMEM with indicated compounds. 48 hours later, 50 uL CTG reagent (Promega) was added to each well and incubated for 15 minutes at room temperature before luminescence measurement. Individual wells of drug-treated (non-DMSO) cells were normalized to the mean of the DMSO-treated wells within an individual experiment. Statistical analysis was performed on the mean of normalized values from individual experiments within treatment groups such that the sample size of each treatment group was equal to the number of independent experiments.

### Seahorse Mitochondrial Stress Test

25,000 cells were seeded in 80 uL full DMEM per well and placed at room temperature for one hour to adhere before being placed in 37C incubator for another two hours. Assay media was made using Seahorse DMEM supplemented to a final concentration of 25 mM glucose and 4 mM glutamine and pH was adjusted to 7.4 after warming to 37C. Inhibitors were diluted in assay media at the following concentrations: 10x oligomycin (15 uM), 10x CCCP (20 uM), 10x antimycin A/rotenone (10 uM each). Before loading cells, all wells had seeding media aspirated and replenished with 180 uL Seahorse media. Reserve oxygen consumption rate was calculated by subtracting the average resting OCR from the average CCCP-treated OCR.

### NAD/NADH-Glo Assay

50,000 cells were seeded per well in 6-well plates in 2 mL full DMEM. After 24 hours, cells were rinsed in PBS, trypsinized, and counted. 5000 cells per condition were assayed per well in white-walled 96-well plates using the Promega NAD/NADH-Glo kit. Cells of independent experiments were seeded and assayed on separate days.

### Cell Accumulation Assay

10,000 cells were seeded in 6-well wells in full DMEM in two plates per independent experiment for day 2 and day 4 cell counts. 24 hours after seeding, media was replaced to fresh DMEM with DMSO or indicated compounds (oligomycin = 2×10-^15^ moles/cell [10 nM], CCCP = 5×10^-14^ moles/cell [250 nM], rotenone = 5×10^-14^ moles/cell [250 nM]). Cells were trypsinized, harvested, and counted using a hemacytometer on day 2 and day 4. Cells of independent experiments were seeded and assayed on separate days.

### Mitochondrial Isolation

Mitochondrial isolation was performed as previously described (Clayton and Shadel, 2014). Cells were cultured in 15 cm plates in full DMEM to full confluence and then trypsinized, harvested, and rinsed once in PBS. The cell pellet was resuspended in 900 uL RSB Hypo Buffer and placed on ice for 15 minutes to allow cells to swell. Cells were transferred to a 5 mL Dounce homogenizer and homogenized with 20 strokes. Cells were transferred to a new 1.5 mL Eppendorf and 600 uL 2.5X MS homogenization buffer was added and mixed. Cells were centrifuged at 1300g for 5 min at 4C and the supernatant containing the mitochondrial fraction was transferred to a new tube. The mitochondrial fraction was centrifuged and transferred two more times to remove remaining nuclei or cells. Mitochondria were pelleted by centrifugation at 15000g for 15 minutes, rinsed in 1 mL of 1X MS homogenization buffer, and pelleted again at 15000g for 15 minutes. Mitochondria were resuspended in buffer (150 mM sodium acetate, 30 mM HEPES, 1 mM EDTA, 12% glycerol (w/v), pH 7.5) at 2 ug protein/uL for further analysis. An aliquot of each lysate was reserved for immunoblot analysis of equal gel loading. Cells of independent experiments were seeded, harvested, and fractionated on separate days.

### Mitochondrial Native PAGE and ETC Complex I In-Gel Activity (IGA) Assay

High resolution clear native (hrCN) PAGE was performed as previously described (Beutner and Jr, 2021). Briefly, isolated mitochondria were solubilized with 2 ug dodecyl maltoside per ug protein on ice for 30 minutes. Solubilized mitochondria were centrifuged at 17,000xg for 10 minutes at 4C. 25 ug solubilized mitochondria was mixed with 1 uL loading buffer (50% glycerol, 0.1% ponceau S) per 10 uL sample. 4-10% acrylamide gradient gels were cast by hand by laying down 1 mL layers at concentrations including and between 4% and 10%. Gels were run at 4C for 1 hour at 100V and then 1 hour at 200V. After electrophoresis, gels for complex I activity assay were placed in complex I IGA buffer (5 mM Tris pH 7.4, 2.5 mg/mL MTT, 0.1 mg/mL NADH) on a shaker for 15 minutes at room temperature. After 15 minutes, buffer was removed, and gel was incubated in 10% acetic acid for 15 minutes on a shaker and then imaged. Complex I IGA band intensities were quantified using FIJI. After gel electrophoresis, gels for Coomassie stain were placed in 0.025% Coomassie G250 in 10% acetic acid at room temperature for 1 hour. Gels were then rinsed overnight in 10% acetic acid in containers with Kimwipes and imaged.

### Mitochondrial DNA Measurement

Measurement of the ratio of mitochondrial to nuclear DNA was performed as previously described (Quiros *et al*., 2017). Cells were cultured in 10-cm dishes and harvested by trypsinization. Cell pellets were lysed in 600 uL buffer (100 mM NaCl, 10 mM EDTA, 0.5% SDS, 20 mM Tris-HCl pH 7.4, 200 ug/mL proteinase K) at 55C for 3 hrs. DNA was precipitated with 250 uL 7.5M ammonium acetate and 600 uL 70% isopropanol and centrifuged at 4C for 10 minutes at 15,000g. The pellet was rinsed with 70% ethanol and centrifuged again and DNA was resuspended in TE buffer. DNA concentration was measured on a NanoDrop and DNA was diluted to 10 ng/uL for qPCR. qPCR was performed in triplicate for each sample and each primer set in 96-well plates with each well containing 10 uL 2x SYBR green qPCR mix, 8 uL of 10 ng/uL DNA, and 2 uL of 10 uM combined forward and reverse primers. The qPCR program run was 95C (5 min), 45 cycles of 95C (10s), 60C (10s), and 72C (20s), then a melting curve. Copy numbers of mitochondrial:nuclear DNA were calculated by the ΔΔCt method as #copies mtDNA = 2*2^(Ct(HK2)-Ct(ND1 or 16S)). Primer sequences can be found in table S3. Student’s T-test was used for statistical analysis.

### Transmission Electron Microscopy

Cultured cells were harvested by trypsinization, rinsed in PBS, and pelleted. Cell pellets were fixed in 1 mL of 4% formaldehyde 0.1M sodium cacodylate at 4C for 1-3 days. Fixative was removed and pellets were rinsed twice in 1 mL 0.1M sodium cacodylate. After the second rinse, cells were placed in osmium tetroxide in 0.1M sodium cacodylate for 45 minutes at room temperature shaking in the dark. Cell pellets were then rinsed once in 0.1M sodium cacodylate and dehydrated by successive 10-minute incubations in 30%, 50%, 70%, 95%, and 100% ethanol. Dehydrated samples were embedded in increasing concentrations (25%, 50%, 75%, 100%) of resin and placed in the oven. Embedded samples were then sectioned and stained with uranyl acetate and lead. Samples were imaged at 30k magnification. Images were analyzed in a randomized and blinded fashion.

### Mitotracker Red CMXRos Staining and Imaging

Cells were seeded on coverslips in 6-well plates at 100,000 cells per well in full DMEM and adhered overnight. To stain, media was replaced with fresh full DMEM + 100 nM Mitotracker Red CMXRos for 45 minutes in a 37C incubator. After staining, media was aspirated and replaced with fresh DMEM for stain washout and cells were placed back in incubator for 60 minutes. Cells were rinsed once in PBS and fixed in 4% formaldehyde in PBS for 15 minutes at RT in the dark. Fixative was removed and nuclei were stained with DAPI-containing PBS for 10 minutes on a shaker in the dark at room temperature. Cells were rinsed twice with PBS on a shaker for 5 minutes each before mounting on slides in Prolong Gold. Slides were imaged on a Zeiss LSM710 and images were exported to TIFFs using ZEN Black software.

### Plasmid Construction and Viral Transduction Cell Line Generation

#### Drp1 Plasmids

Mouse Drp1 was PCR amplified from pcDNA3.1 mDrp1 (Addgene #34706) and cloned into pLenti BlastR using InFusion cloning. Drp1 mutants were generated using InFusion cloning.

#### CRISPR

plentiCRISPRv2 (Addgene #52961) was digested with BsmBI and annealed sgRNA oligos were ligated with T4 ligase. For double CRISPR cells, the puromycin resistance transgene from plentiCRISPRv2 was swapped for a neomycin resistance transgene to generate plentiCRISPRv2 NeoR and sgRNAs were cloned in using the same protocol. Lentivirus was generated in 293T cells by calcium phosphate cotransfection with psPAX2 (Addgene #12260), pCMV-VSV-g (Addgene #8454), and plentiCRISPRv2 vectors. Media was removed from 293T cells 24 and 48 hours after transfection, filtered through 0.45 um PES filters, and applied to target cells with polybrene (5 ug/mL). Infected cells were selected in full DMEM with puromycin (3 ug/mL for mouse cells, 1 ug/mL for A549) or neomycin (1 mg/mL mouse and A549). CRISPR guide RNA sequences can be found in Table S2.

### Immunoblotting

Cell pellets were lysed on ice in RIPA buffer with protease/phosphatase inhibitor cocktail (Roche). Protein concentration was determined by BioRad Protein Assay (BioRad) and 20 ug protein was resolved by SDS-PAGE in 10% acrylamide gels, transferred to PVDF membrane (Millipore), blocked in 5% fat-free milk, and probed with indicated antibodies.

### TMRE Flow Cytometry

500,000 cells were seeded in 10 mL full DMEM in 10-cm plates and adhered overnight. The next day, media was changed to full DMEM with DMSO, oligomycin, or CCCP at equimolar amounts per cell as the cellular accumulation experiment using these same compounds (oligomycin = 2×10-^15^ moles/cell [100 nM], CCCP = 5×10^-14^ moles/cell [2.5 uM]) and incubated for one hour at 37C. Cells were trypsinized, counted, and 250,000 per treatment were incubated in 500 uL PBS + 4 mM glutamine + 25 mM glucose + 100 nM TMRE for 30 minutes at 37C before analysis by flow cytometry on an Attune NxT flow cytometer. Data were analyzed in FCS Express. TMRE intensity was determined in cells gated solely on the live cell population as determined by SSC x FSC.

## QUANTIFICATION AND STATISTICAL ANALYSIS

Statistical analysis and data presentation were performed using GraphPad Prism v7. Statistical tests used and statistical significance are indicated in figure panels and legends.

## Resources

| REAGENT or RESOURCE | SOURCE | IDENTIFIER |
| --- | --- | --- |
| **Antibodies** |  |  |
| Mouse monoclonal anti-Opa1 (WB) | BD Biosciences | 612606 |
| Rabbit polyclonal anti-Opa1 (IHC) | GeneTex | 48589 |
| Rabbit monoclonal anti-Drp1 (WB + IHC) | Cell Signaling Tech. | 5391 |
| Rabbit monoclonal anti-beta tubulin | Cell Signaling Tech. | 2128 |
| Rabbit monoclonal anti-beta Actin | Cell Signaling Tech. | 8457 |
| Rabbit monoclonal anti-PARP | Cell Signaling Tech. | 9542 |
| Rabbit monoclonal anti-SDHA | Cell Signaling Tech. | 11998 |
| Rabbit monoclonal anti-VDAC | Cell Signaling Tech. | 4866 |
| **Bacterial and virus strains** |  |  |
| Ad5-CMV-Cre | Baylor Vector Core | N/A |
| Ad5-CMV-Empty | Baylor Vector Core | N/A |
| **Chemicals, peptides, and recombinant proteins** |  |  |
| MYLS22 Opa1 inhibitor | MedChemExpress | HY-136446 |
| Avertin (2-2-2,-tribromoethanol) | Fisher | AAA1870606 |
| **Critical commercial assays** |  |  |
| NAD/NADH-Glo | Promega | G9071 |
| Seahorse Mitochondrial Stress Test | Agilent | 103015-100 |
| CellTiter-Glo | Promega | G7570 |
| **Experimental models: Cell lines** |  |  |
| A549 human lung adenocarcinoma | ATCC | CCL-185 |
| KP, KPD, KPO, KPDO mouse lung adenocarcinoma | This study | N/A |
| **Experimental models: Organisms/strains** |  |  |
| Mouse: KP: *KRAS*<sup>LSL-G12D/WT</sup>; *TP53*<sup>FL/FL</sup> | DuPage et al., 2009 | N/A |
| Mouse: D: *DRP1*<sup>FL/FL</sup> | Wakabayashi et al., 2009 | N/A |
| Mouse: O: *OPA1*<sup>FL/FL</sup> | Zhang et al., 2011 | N/A |
| **Recombinant DNA** |  |  |
| plentiCRISPRv2 | Addgene | 52961 |
| plenti BlastR Luciferase-V5 | Addgene | 21474 |
| pcDNA3.1 mDrp1 | Addgene | 34706 |
| **Software and algorithms** |  |  |
| Graphpad Prism v8.3 | GraphPad | N/A |
| Fiji | ImageJ | https://imagej.net/software/fiji/downloads |
| QuPath v0.3.0 | Bankhead et al., 2017 | https://qupath.github.io/ |
| Aperio ImageScope | Leica Biosystems | N/A |
| FCS Express | DeNovo Software | N/A |
| ZEN Black | Zeiss | N/A |

---

**Table S1:**
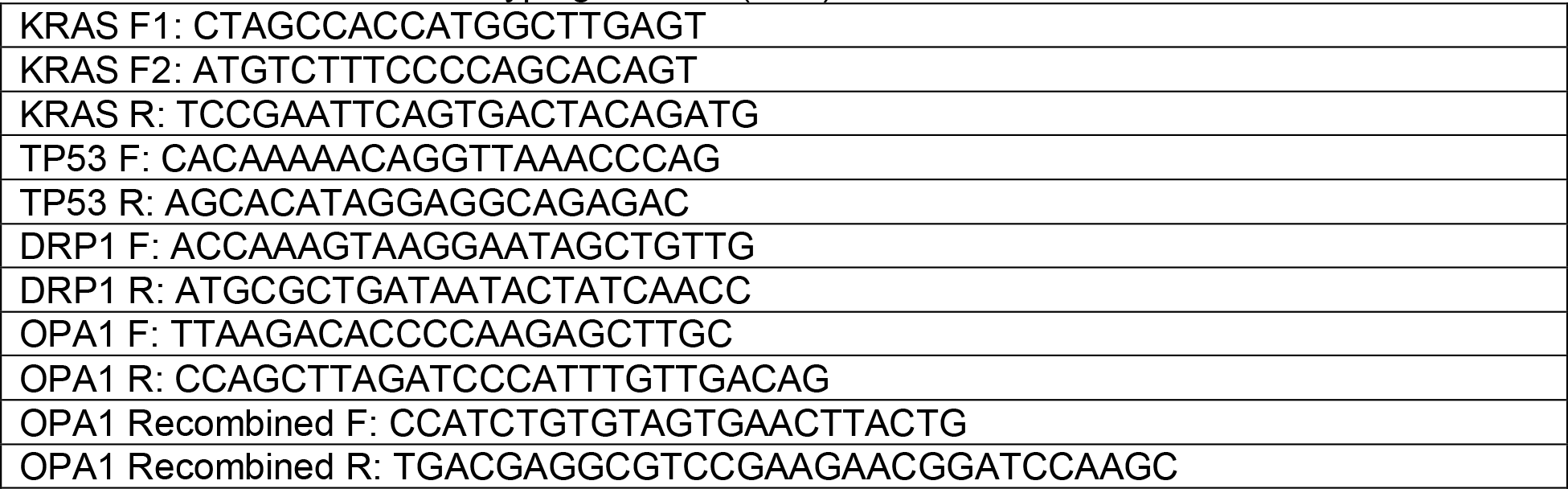
KPDO Mouse Genotyping Primers (5’-3’)

**Table S2:**
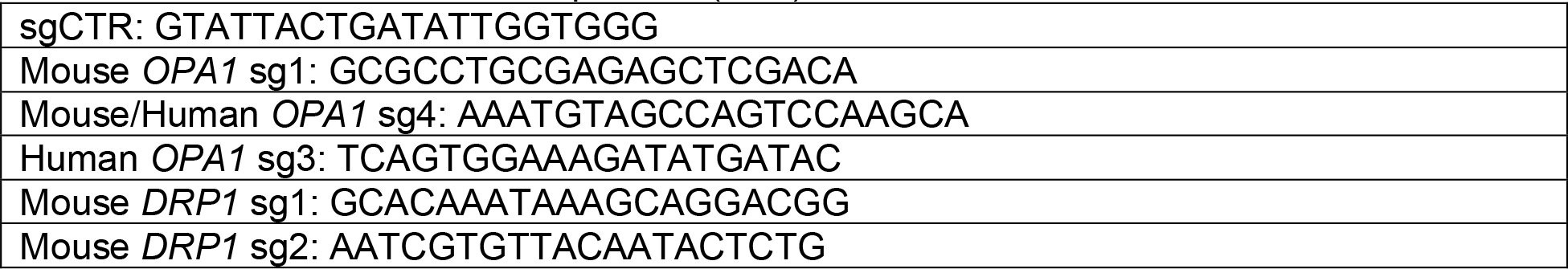
CRISPR Guide RNA Sequences (5’-3’)

**Table S3:**
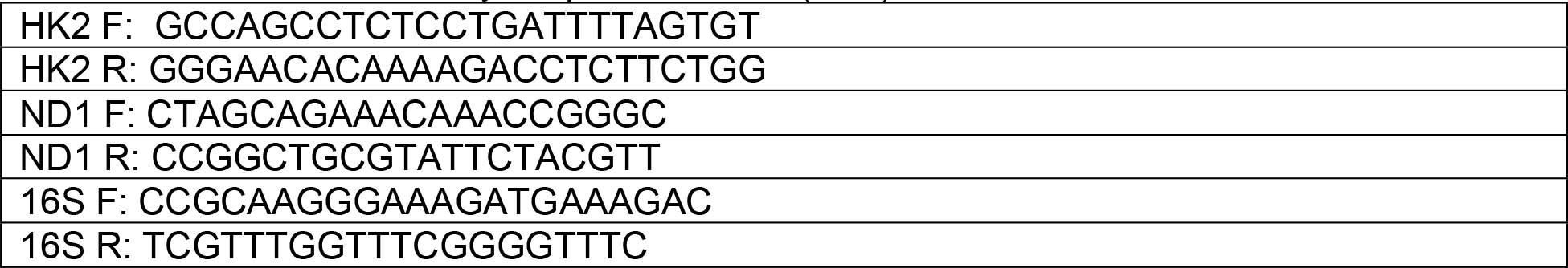
mtDNA:nDNA Analysis qPCR Primers (5’-3’)

**Figure S1:**
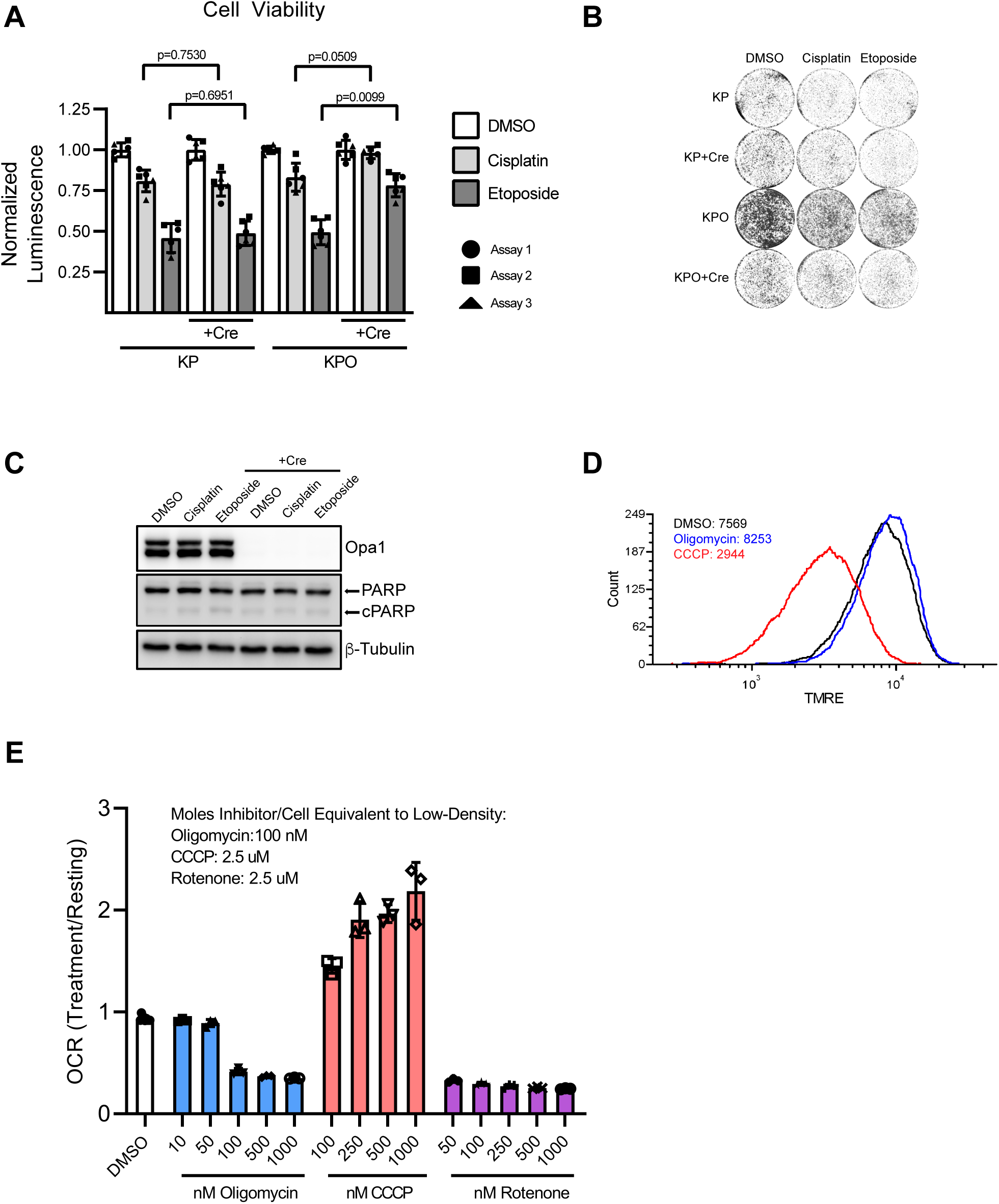
Opa1 deletion does not universally sensitize LUAD tumor cells to apoptosis. A. CellTiter-Glo measurement of cell viability of uninfected or AdCre-infected KP and KPO tumor cells treated with DMSO, cisplatin (2 uM), or etoposide (2 uM) for 48 hours. All technical replicates from independent experiments are shown. Individual wells of drug-treated (non-DMSO) cells were normalized to the mean of the DMSO-treated wells within an individual experiment. Statistical analysis was performed on the mean of normalized values from individual experiments within treatment groups such that the sample size of each treatment group was equal to the number of independent experiments. n=3 independent experiments. Bars represent mean±SD. Student’s T-test. B. Crystal violet staining of uninfected or AdCre-infected KP and KPO tumor cells treated with DMSO, cisplatin (2 uM), or etoposide (2 uM) for 48 hours. Cells were seeded at 10,000 cells/12-well in 1 mL 10% FBS DMEM. C. Immunoblot of Opa1 and PARP expression in uninfected or AdCre-infected KPO tumor cells treated with DMSO, cisplatin (2 uM), or etoposide (2 uM) for 48 hours. D. TMRE flow cytometry of uninfected KPO tumor cells treated with equal number of moles oligomycin/CCCP per cell as the cellular accumulation experiment in Figure 4F. Cells were gated solely on live cell population based on FSCxSSC. Values indicate TMRE median fluorescence intensity per group. n>20,000 cells per treatment group. E. Treatment/resting oxygen consumption rate of uninfected KPO tumor cells with indicated concentrations of compounds measured by Seahorse bioanalyzer. n=3 wells per condition. Bars represent mean±SD.

**Figure S2:**
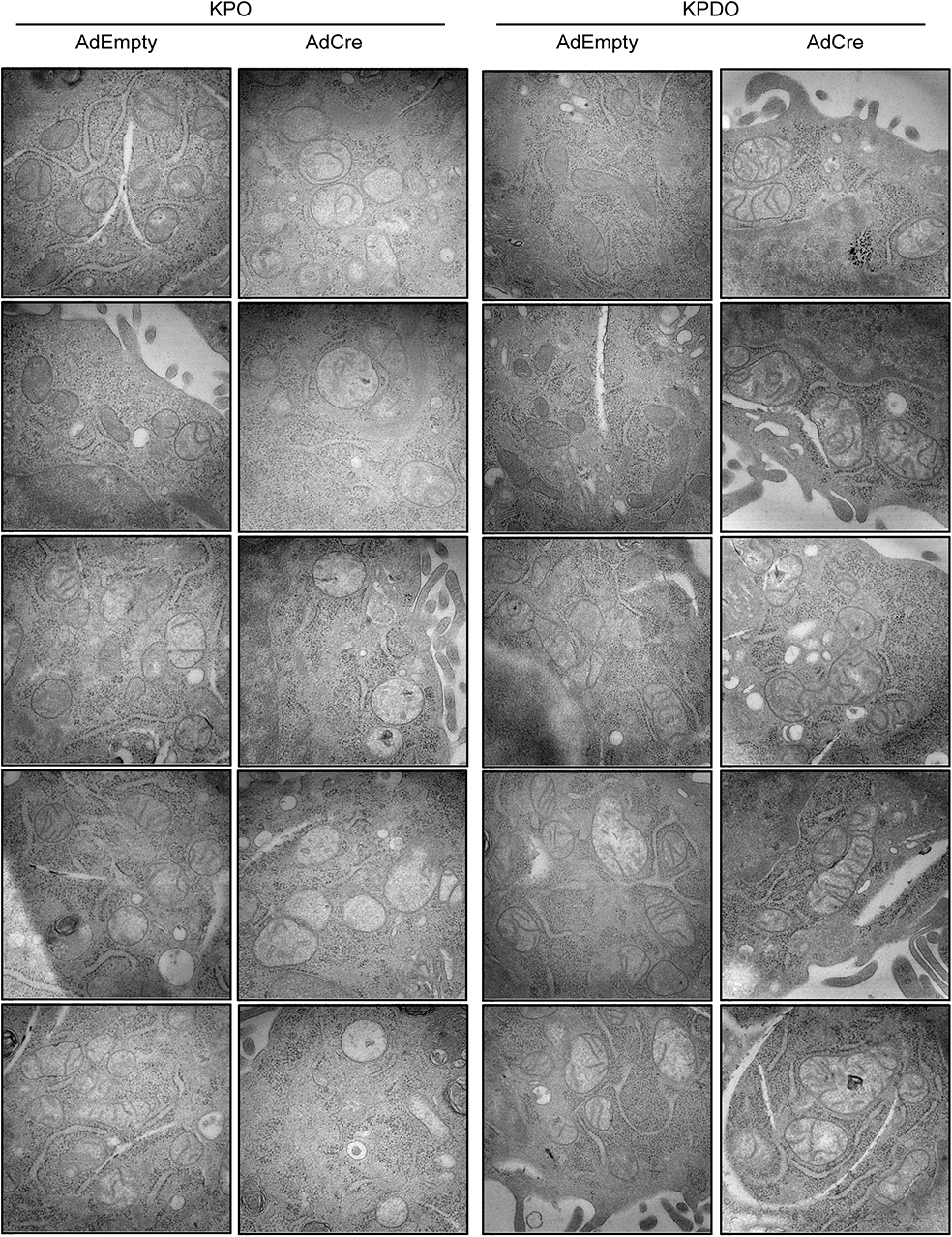
Opa1 deletion impairs mitochondrial cristae morphology in a Drp1-dependent manner. Representative TEM images of mitochondria and cristae morphology in AdEmpty- or AdCre-infected KPO and KPDO tumor cells. Magnification = 30k. Scale bar. = 200 nm.

## Notes

### Competing Interest Statement

The authors have declared no competing interest.

